# Defining the cellular and molecular features of nerve-invaded cancer cells using a newly characterized experimental model

**DOI:** 10.1101/2025.03.21.644644

**Authors:** Nickson Joseph, Sanskriti Shrestha, Ivanka Sassmannshausen, Sandra Mizkus, Paulos Chumala, Bhadrapriya Sivakumar, Viswanath Baiju, Shahid Ahmed, Henrike Rees, George S. Katselis, Anand Krishnan

## Abstract

Perineural invasion (PNI) is the invasion of cancer cells into nerves. Although PNI is a risk factor for cancer recurrence and metastasis, the lack of *in vitro* experimental models representing natural PNI challenges basic studies and therapeutic screening. In this work, we fully characterized a dorsal root ganglia (DRG)-nerve explant model for PNI and demonstrated the characteristic cellular and molecular features of cancer cells undergoing natural PNI. Briefly, thoracic and lumbar DRGs intactly connected to nerves were co-cultured with breast and prostate cancer cells in a 3D matrix for two weeks. Time-dependent brightfield and fluorescence imaging captured the complex interactions of cancer cells, neurons, axons, and Schwann cells within nerves in the DRG-nerve explant, demonstrating the natural invasion of cancer cells. Fundamental investigations showed that the autonomic neurotransmitters norepinephrine and acetylcholine significantly promote PNI. We also demonstrated increased survival of PNI cells against the cytotoxic drug cisplatin. Additionally, we characterized the proteomics profile of PNI cells for future theranostics applications and validated the results using patient breast tumor samples. Overall, this work characterized and established a clinically relevant model for PNI and revealed the cellular crosstalk of PNI cells within nerves. The established model is suitable for fundamental studies and therapeutic screening pertaining to PNI.

## Introduction

Perineural invasion (PNI) is the invasion of cancer cells into peripheral nerves and is a known risk factor for cancer metastasis and recurrence ^1–4^. The association between PNI and the development of metastasis and disease recurrence has been shown in breast, prostate, liver, head and neck, colorectal and pancreatic cancers ^2, 4–8^. The clinical diagnosis of PNI includes microscopic evaluation of Hematoxylin and Eosin (H&E) stained tumor sections by an anatomical pathologist focusing on nerve infiltration, and cancer cell distribution in and around nerves. While imaging methods, such as, MRI, CT, and PET are useful for clinical diagnosis of PNI, they are not the best in providing accurate information, such as, how far from the primary tumor the cancer cells have invaded the nerves, especially the internal layers of nerves ^9^. Uncovering such deeper level information will assist the oncology team in accurately predicting disease prognosis and designing treatment plans. However, the limited knowledge of the molecular signature of PNI-competent cells, herein referred to as PNI cells, challenges the development of molecular approaches targeted to enhancing the visualization of PNI cells. Revealing the molecular signature of PNI cells should also facilitate the development of targeted therapies for PNI.

A pre-clinical model representing true characteristics of PNI is currently unavailable and is challenging to develop. Animal tumor models that are currently available are short-term models meeting ethical requirements, while natural PNI may take a longer time to develop. Although, several *in vitro* models exist, they either fail to induce natural PNI or lack intact nerve preparations, failing to match the milieu requirements for natural PNI^13^. This highlights the demand for a better *in vitro* model containing intact nerve preparations, enabling natural PNI, for PNI-related molecular studies. Interestingly, Ayala et al., in the early 2000 showed that co-cultures of Dorsal Root Ganglia (DRG)-nerve explant and cancer cells induce a mutual growth response^10^. However, a systematic characterization of this model for true PNI, the invasion of cancer cell into nerve layers, and the presence of healthy axons and supporting Schwann Cells (SCs), the basic components of nerves, within nerves, has not been done. Here, we fully characterized the DRG-nerve explant and cancer cells co-culture model and demonstrated that this model recapitulates natural PNI and comprises of nerve-resident healthy axons and SCs. We also characterized the molecular signature of PNI-competent breast cancer cells using this model and validated unique markers. We demonstrated that this model is useful for exploring the fundamental biology of PNI and screening potential anti-PNI therapies. Moreover, the molecular markers of PNI cells identified in this work should facilitate the development of precise molecular diagnostics and therapeutic targets for PNI.

## Materials and Methods

### Cell culture and chemicals

MDA MB-231, LNCaP, and LASCPC-01 cells were purchased from the American Type Culture Collection (ATCC, USA). MDA MB-231 cultures were maintained in DMEM/F12 media (Life Technologies:11-330-057) containing 10% FBS (Life Technologies:12483020) and 1% antibiotic-antimycotic cocktail (Cytvia HyClone:SV3007901). LNCaP cultures were maintained in RPMI (Cytvia HyClone:SH3002701), 10% FBS and 1% antibiotic-antimycotic cocktail. LASCPC-01 cultures were maintained in RPMI, 10% FBS, 1% insulin transferrin selenium (Gibco™: 41400045), 10nM beta-estradiol (Sigma-Aldrich: E2758), 10nM hydrocortisone (Sigma-Aldrich:H0888) and 1% antibiotic-antimycotic cocktail. All cultures were maintained in the incubator at 37°C and 5% CO_2_. Cisplatin (232120), norepinephrine (A9512) and acetylcholine (A2661) are purchased from Sigma-Aldrich. Primary human breast tumor samples were purchased from the Alberta Cancer Research Biobank (ACRB).

### Adenoviral transduction

For generating GFP^+^ cancer cells, the cells were transduced with adenovirus expressing enhanced green fluorescent protein under the control of a CMV promoter (Ad-GFP; Vector Biolabs:1060). Briefly, MDA-MB-231, LASCPC 01 or LNCaP cells were seeded in T25 culture flasks containing the standard culture media. After 24 hours, Ad-GFP (100 MOI) along with 8 µg/mL Polybrene (Sigma-Aldrich:TR-1003) was added to the culture for adenoviral transduction. After 24 h, the media was replaced with fresh media. Fluorescence images of GFP^+^ cells were captured thereafter to ensure successful transduction.

### *In vitro* model of PNI

Thoracic and/or lumbar dorsal root ganglia (DRG) with ∼3-4 mm nerve segments attached to them were isolated from 4-6 weeks old male SD rats. The DRG-nerve preparations were then embedded in Cultrex extracellular matrix (R & D Systems:3433-010-01) in a 24 well plate with a single DRG-nerve preparation embedded per well. Thereafter, 10,000 regular or GFP^+^ cancer cells were seeded at three different spots closer to the nerve endings of the DRG-nerve preparation. The co-cultures were maintained in DMEM/F12 containing 10% FBS, 0.1% N2 (ThermoFisher Scientific:17502048), 0.1% BSA (HyClone:SH3057402), 100 ng/mL NGF (ThermoFisher Scientific:13257-019), and 1% antibiotic/antimycotic cocktail at 37°C and 5% CO_2_ for two weeks, with the media replaced with fresh media every two days. Brightfield and fluorescent images of the preparation were captured at regular intervals using Axio Observer inverted microscope (Zeiss). After two weeks, the DRG-nerve preparations were harvested from the matrix for immunohistochemistry and cell sorting experiments.

### Immunohistochemistry

DRG-nerve preparations were washed in PBS thrice for clearing the Cultrex matrix. The preparations were then incubated in Zamboni’s buffer (2% formaldehyde (Fisher Scientific: F79-500) and 0.5% picric acid (Ricca Chemical:5860-16) in PBS) overnight at 4 °C. The next day, the tissues were washed thrice with PBS and incubated in 20% sucrose (Fisher Scientific: BP220-1) overnight at 4 °C. The tissue blocks were then made using optimal cutting temperature compound (OCT, Fisher Scientific:23-730-571) and the blocks were stored at −80°C until further use.

For immunohistochemistry work, 12 µm thick sections of the DRG-nerve preparations were made using a cryostat. The sections were blocked using 5% donkey serum (Fisher Scientific:56-646) and permeabilized using 0.3% Triton X (Fisher Scientific: BP151-100). The sections were then washed using PBS and incubated with primary antibodies for 1 h. The primary antibodies used were βIII tubulin, (Sigma-Aldrich: MAB1637, dilution:1:500), NF200 (Sigma-Aldrich: N0142, dilution:1:500), GFAP (Invitrogen:PA110004, dilution:1:500), cytokeratin (Novus Biologicals: NB120-6401, dilution:1:200), and Ki67 (Invitrogen: PA5-16785, 1:100). For PNI marker validation experiments, hemopexin (Invitrogen:MA5-32799, dilution:1:200) was used. The slides were then washed thrice with PBS followed by secondary antibody incubation was done for 40 minutes. The secondary antibodies used were, anti-chicken Alexa Fluor 647 (Life Technologies: A-21449, dilution:1:100), anti-mouse Alexa Fluor 488 (Invitrogen: A-11001, dilution:1:100), anti-mouse Alexa Fluor 546 (Invitrogen: A21045, dilution:1:100), anti-mouse Alexa Fluor 647 (Invitrogen: A-21235, dilution:1:100) and anti-rabbit Alexa Fluor 647 (Invitrogen:A-21245, dilution:1:100). The slides were then washed thrice with PBS followed by mounted using SlowFade^TM^ Diamond Antifade mountant with DAPI (Life Technologies: S36973).

For experiments designed to visualize only GFP^+^ cancer cells in sections, the slides with DRG-nerve sections were washed thrice in PBS and directly mounted. The PNI cells were then manually counted from the images taken at 20x magnification. For quantification of PNI cells, several images representing multiple fields/samples were generated and the average number of PNI cells/field is tabulated.

### Hematoxylin and Eosin (H&E) Staining

A total of 98 human breast tumor samples were stained using H&E. Samples were initially fixed using Zamponi’s buffer overnight at 4°C, washed thrice in PBS, and then incubated in 20% sucrose solution overnight at 4°C. The tissue blocks were made using OCT compound, and then, 12 µm sections were made using a cryostat. Each slide contained 3 sections at three different depth levels (∼48 µm apart) of the sample. The slides were passed through 95% of ethanol and distilled water for initial hydration and then stained using Modified Harris hematoxylin solution (Sigma-Aldrich: HHS16) for nuclear staining. Slides were then rinsed in tap water and again washed in distilled water to adjust the intensity of hematoxylin stain. Following the washes, sections were stained using Eosin Y stain (Fisher Scientific: E511-25) for cytoplasmic staining. The slides were then passed through a series of alcohol changes (twice in 95% ethanol and twice in absolute ethanol) for dehydration and rinsed twice in xylene for clearing the samples. Finally, the sections were mounted using the DPX mountant (Sigma-Aldrich:06522). The stained sections were evaluated for PNI incidence by a pathologist.

### qRT-PCR

2 µg of total RNA was converted into cDNA using a cDNA synthesis kit (Applied Biosystems:4368814) as per the manufacturer’s instructions. The primers used were: CPLANE1 (5’-ACCTTGGCCACATGGAACTG-3’ (F) and 5’-GCAGACTATGACACGGAGCA-3’ (R); hemopexin (5’-AGCAGTGGATGCTGCCTTTTCC-3’ (F) and 5’-TTCTCCAGCCGCTTCGGATAAC-3’ (R); beta-actin (5’-CACCATTGGCAATGAGCGGTTC-3’ (F) and 5’-AGGTCTTTGCGGATGTCCACGT-3’ (R). PowerUp™ SYBR™ Green Master Mix (Applied Biosystems: A25776) was used for cDNA amplifications and the reactions were carried out in a QuantStudio™ 3 Real-Time PCR System (Applied Biosystems).

### Cell sorting

GFP^+^ cancer cells were sorted using a flow cytometer (FACSMelody™ Cell Sorter, BD Biosciences). Individual cell suspension was initially made from the DRG-nerve preparations from the *in vitro* PNI model. Briefly, the DRG-nerve preparations were enzymatically digested using 0.1% collagenase (Life Technologies:17104019) for 90 minutes at 37°C. After enzymatic digestion, individual cells were mechanically dissociated using repeated pipetting. The suspension was then laid over 15% BSA and centrifuged at 800 rpm for 6 minutes to clear the debris that accumulate in the supernatant. The pellet containing individual cells was then washed in PBS and suspended in PBS containing 2% FBS for sorting GFP^+^ cells using a flow cytometer. The GFP^+^ cancer cells were pooled from ∼50 DRG-nerve preparations for each of the two biological replica yielding sufficient amount of protein (>12 µg/sample) for the proteomics studies.

### Proteins In-Solution Tryptic Digestion

Each protein sample was diluted with 5 µL of 100 mM ammonium bicarbonate (ABC) buffer (Fisher Scientific) and 50 µL trifluroethanol (TFE, Fisher Scientific). The samples were then treated with 1 µL of 1 M dithiothreitol (DTT, MP Biomedicals) while shaking at 300 RPM at 60°C for 60 min. The samples were then alkylated with 100 µL of 110 mM iodoacetamide (IAA, Fisher Scientific) at 37°C for 30 min and dried in a speed-vac. Proteins in the samples were treated with 1 mL cold acetone followed by refrigeration at −80°C for 60 min. The acetone was then carefully removed after centrifugation. Samples devoid of interferences were dried in a speed-vac and a buffer-containing trypsin (Promega Corporation) solution (50 ng/µL trypsin in 1 mM HCl / 100 mM ABC) was added to the samples in a 40:1 protein:trypsin ratio. The samples were incubated in a shaker at 300 RPM overnight at 37°C. Trypsin buffer at the same ratio was added again in the morning to ensure complete digestion of proteins into peptides. After 2 h of further incubation at 37°C, digested peptides were dried in speed-vac and stored at −80°C until further analysis.

### Mass Spectrometry Workflow

Protein digested samples were reconstituted in 20 µL of MS grade water:ACN:FA (97:3:0.1 v/v) followed by vortexing for 1 to 2 min to achieve peptide solubility. The resulted solutions were centrifuged at 18000 g for 10 min at 4°C. A 15 µL aliquot of each sample was transferred to a mass spectrometry vial for liquid chromatography-tandem mass spectrometry (LC-MS/MS) analysis. All mass spectral analyses were performed on an Agilent 6550 iFunnel quadrupole time-of-flight (QTOF) mass spectrometer equipped with an Agilent 1260 series liquid chromatography instrument and an Agilent Chip Cube LC-MS interface (Agilent Technologies). Chromatographic peptide separation was accomplished using a high-capacity Agilent HPLC-Chip II: G4240-62030 Polaris-HR-Chip 3C18 consisting of a 360 nL enrichment column and a 75 µm × 150 mm analytical column, both packed with a Polaris C18-A, 180Å, 3 µm stationary phase. Samples were loaded onto the enrichment column with solvent A (0.1% formic acid in water) at a flow rate of 2.0 µL/min. Samples loaded to enrichment column were transferred onto the analytical column, and peptides were separated with a linear gradient solvent system consisted of solvent A and solvent B (0.1% formic acid in acetonitrile). The composition of the linear gradient during the chromatographic analysis was 3–25% solvent B for 105 min, 25-40% solvent B for 15 min, and then 40–90% solvent B for 5 min at a flow rate of 0.3 µL/min. Positive-ion electrospray mass spectral data were acquired using a capillary voltage set at 1900 V, the ion fragmentor set at 360 V, and the drying gas (nitrogen) set at 225°C with a flow rate of 12.0 L/min. Spectral results were collected over a mass range of 250–1700 (mass/charge; m/z) at a scan rate of 8 spectra/s. MS/MS data were collected over a range of 100–1700 m/z and a set isolation width of 1.3 atomic mass units. The top 20 most intense precursor ions for each MS scan were selected for tandem MS with active exclusion for 0.25 min.

### Protein Identification

The acquired spectral data were searched against the SwissProt *Homo sapiens* database (March 2024 UniProt release), using Spectrum Mill (Agilent Technologies Canada Ltd., Canada) as the database search engine. Search parameters included a fragment mass error of 50 ppm, a parent mass error of 20 ppm, trypsin cleavage specificity, and carbamidomethyl as a fixed modification of cysteine. In addition, the following variable modifications were used: oxidized methionine, carbamylated lysine, acetyl lysine, pyroglutamic acid, deamidated asparagine, phosphorylated serine, threonine, and tyrosine, semi-trypsin non-specific C-terminus and semi-trypsin non-specific N-terminus. Validation of Spectrum Mill results was performed at peptide and protein levels (1% false discovery rate). Protein lists of treated and control groups were exported to the Mass Profiler Professional (MPP, version 15.0, Agilent Technologies) software for statistical analysis. Data analysis was carried out based on the total spectra intensity of the proteins which were considered as entities in MPP. The baseline of the spectra was adjusted to the median across all samples. The entities were then filtered based on their frequency of occurrence across at least all replicates of one treatment. Unpaired T-test was performed, with a p-value of <0.05 considered to be significant. Benjamini-Hochberg corrected p-values were calculated to overcome the problem of multiple test analysis (false discovery). Proteins that showed ≥ 2.0-fold change in expression in PNI cells compared to control (non PNI cells isolated from areas away from the DRG-nerve preparations in the *in vitro* PNI model) were considered differentially regulated proteins and these proteins were considered for functional enrichment analysis. The mass spectrometry proteomics data have been deposited to the ProteomeXchange Consortium via the PRIDE^11, 12^ partner repository with the dataset identifier PXD059147.

### Functional enrichment analysis of differentially regulated proteins and STRING analysis of protein interaction network

Functional annotation and pathway enrichment analyses were performed using the PANTHER database (PANTHER 19.0). UniProt IDs of differentially regulated proteins was uploaded to the PANTHER online tool, selecting the Gene Ontology (GO) terms and Pathway options. Functional classifications were assessed under molecular function and biological pathway categories. Pathway enrichment analysis was conducted using default settings, with the background set for Homo sapiens. Graphs were generated using PANTHER’s visualization tools. Protein-protein interaction networks were constructed using STRING 12.0. The UniProt IDs of differentially regulated proteins were submitted to STRING, with 0.4 as the interaction score threshold. Interaction types considered included known, predicted and other interactions from experimental data, co-expression, curated and text mining databases. Visualization of interaction networks was done using STRING’s network viewer.

### Statistical Analysis

All statistical analyses were performed using the GraphPad Prism 10 Software. Results were presented as mean ± standard error. The specific analysis performed (standard ‘t’ test, ANOVA), including the number of replicas/groups, and the statistical significance achieved, are presented in the corresponding figure legends. p-values of ≤ 0.05 was considered statistically significant.

## Results

### Characterization of the *in vitro* PNI model comprising an intact nerve preparation

An ideal experimental model of PNI should include intact nerve endings as seen in natural tumor microenvironment. To recapitulate a nerve enriched tumor microenvironment favoring PNI, we co-cultured cancer cells and a DRG with ∼3-4 mm intact nerve attachment (DRG-nerve preparation) in Cultrex extracellular matrix for two weeks. The Cultrex provided a nutrient rich matrix for cancer cells to migrate in a 3D environment while the proximity of cancer cells to an intact nerve, as in natural tumor microenvironment, promoted cancer cells’ migration toward nerve endings in a time dependent manner, as evident from bright field images shown in **Figure 1A-C**. We used breast (MDA-MB-231) and prostate (LNCaP and LASCPC-01) cancer cell lines for these experiments, and all three cell lines showed tropism toward the DRG-nerve tissue at basal culture conditions. In a separate experiment, we used GFP^+^ MDA-MB-231 cells to visualize the cancer cell-nerve interface. Axons (elongated projections of neurons) and SCs, the supporting glial cells, are the two major types of cells present in peripheral nerves. From the brightfield images, it appeared that SCs emerging from the nerve endings enrich the immediate external milieu of the nerve and potentially provide directional cues to migrating cancer cells **(Figure 1D, E (top)).** While both axons and SCs can generate elongated processes, the apparent presence of cell bodies along with thicker cytoplasmic extensions suggest that most cells in the immediate external milieu of nerve are SCs. At the same time, we found the presence of GFP^+^ cancer cells in the SC bed moving towards the nerve tip suggesting initiation of nerve invasion **(Figure 1E (bottom))**.

**Figure 1:**
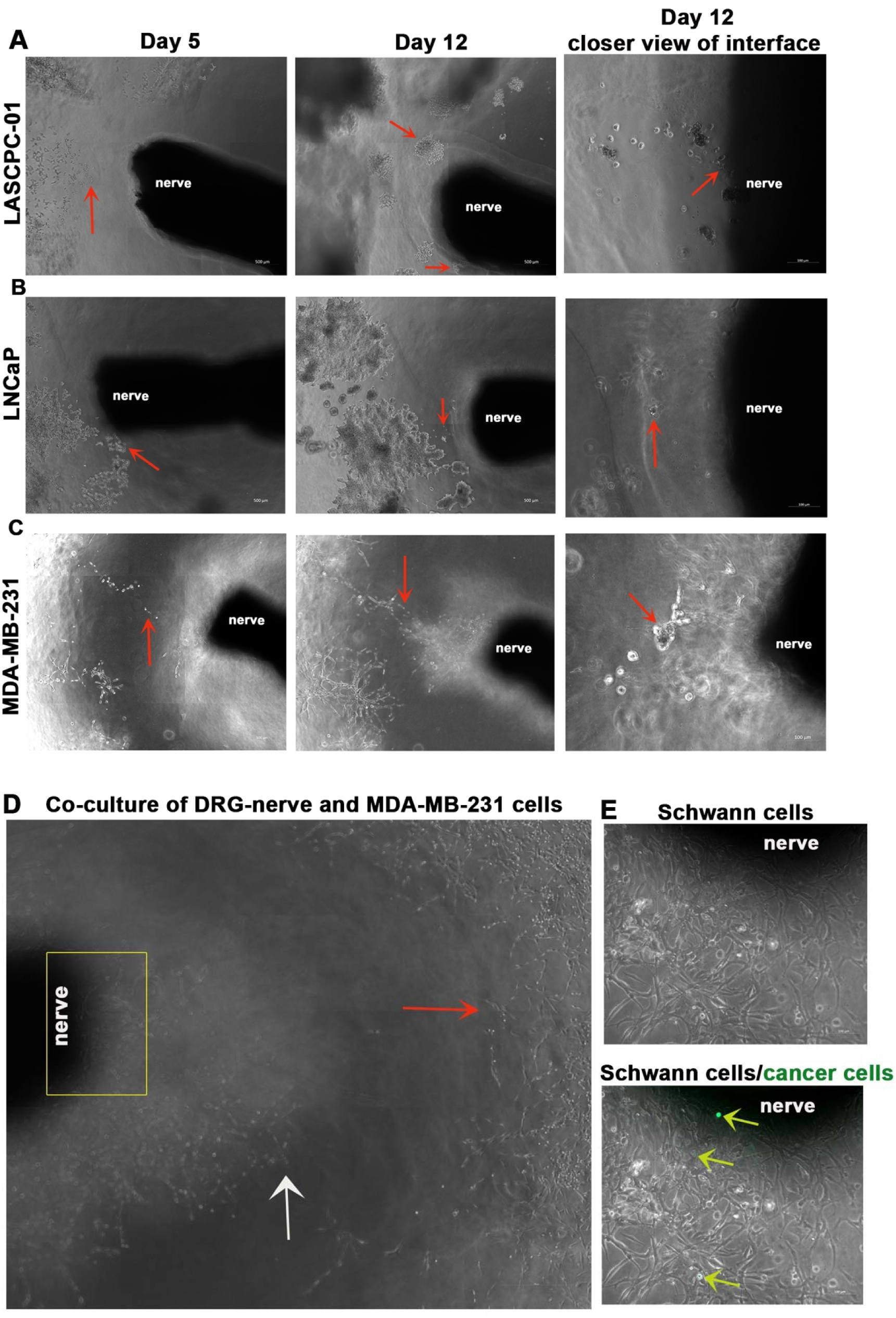
Nerve and cancer cells interact each other in a time dependent manner in the *in vitro* PNI model: **(A-C)** Bright field images show time dependent tropism of cancer cells (red arrows: LASCPC-01, LNCaP, MDA-MB-231) toward nerve endings on days 5 and 12 in the in vitro PNI model (scale bar, 500 µm). A closer view of the nerve-cancer cell interface on day 12 is also shown on the far right (scale bar, 100 µm). **(D)** Nerve-cancer cell interface shows emerging SCs (white arrow) from the nerve endings migrating towards cancer cells (red arrow) (scale bar, 500 µm). **(E-top)** Enlarged view of the boxed region in ‘D’ shows the presence of SCs immediately outside the nerve endings in the in vitro PNI model (scale bar, 100 µm). **(E-bottom)** A combination of brightfield and fluorescent images of the field presented in ‘**E-top’** shows the presence of GFP^+^ cancer cells (yellow arrows) migrating towards the nerve endings along the SC bed (scale bar, 100 µm).

### Cancer cells interact with SCs within the nerve milieu and sensory neurons in the DRGs in the *in vitro* PNI model

We next investigated whether the *in vitro* PNI model (the DRG-nerve-cancer cell co-culture model) allows cancer cell invasion and movement within nerves, as naturally occurs in PNI. To evaluate this, sections of DRG-nerve preparations were examined for the presence of cytokeratin, which marks cancer cells. All three cell lines tested in the *in vitro* model invaded the nerves, as evident from the presence of cytokeratin^+^ cells within nerves, adhering to the strict definition of PNI (**Figure 2A**). To further verify the invasion of cancer cells in this model, we used GFP^+^ MDA-MB-231 cells and examined the presence of GFP^+^ cells (cancer cells) in nerve sections after two weeks. Consistent with the cytokeratin staining experiments, we found GFP^+^ MDA-MB-231 cells within nerves verifying the occurrence of a true PNI in the model (**Figure 2B, C**). We also examined the distribution of GFP^+^ LASCPC-01 cells within nerves in the *in vitro* model and found that they also invade nerve layers (**Figure S1**). It is thus clear from these observations that the DRG-nerve-cancer cell co-culture model established here can mimic natural PNI *in vitro*. We further examined the potential crosstalk of cancer cells with axons and SCs within the nerve tissue to identify which type of cells physically interact and potentially support the survival and migration of cancer cells within the nerve milieu. Axons are marked by staining them with neurofilament 200 (NF200) or βIII tubulin while SCs are marked by staining them with glial fibrillary acidic protein (GFAP). NF200 stains large diameter axons while βIII tubulin stains all types of axons regardless of their size. The sections of DRG-nerve showed intact NF200^+^ strands indicating the presence of intact axons in the 14-day old model (**Figure 2B)**. However, the abundance of NF200^+^ axons was much lower in the PNI model compared to the axon abundance in normal sciatic nerve, indicating that large diameter axons may degrade to some extend or they lose NF200 expression in the *in vitro* model (**Figure S1A**). However, the model showed continuous strands of βIII tubulin^+^ axons within nerves, as in normal sciatic nerve, confirming the presence of intact axons in the model (**Figure 2C, Figure S1A**). We also found high abundance of SCs within the nerves of the PNI model, as in normal sciatic nerve, further verifying the presence of natural nerve constituents in a 14-day old *in vitro* model (**Figure 2B, Figure S1A**). The GFP^+^ MDA-MB-231 cells were found randomly distributed within nerves, while showing physical proximity preferentially to SCs, suggesting that SCs may support PNI cells within the nerve segments (**Figure 2B**). Such closer cellular interactions were also observed between prostate cancer cells and SCs, substantiating that SCs support cancer cells within nerves (**Figure S1B**).

**Figure 2:**
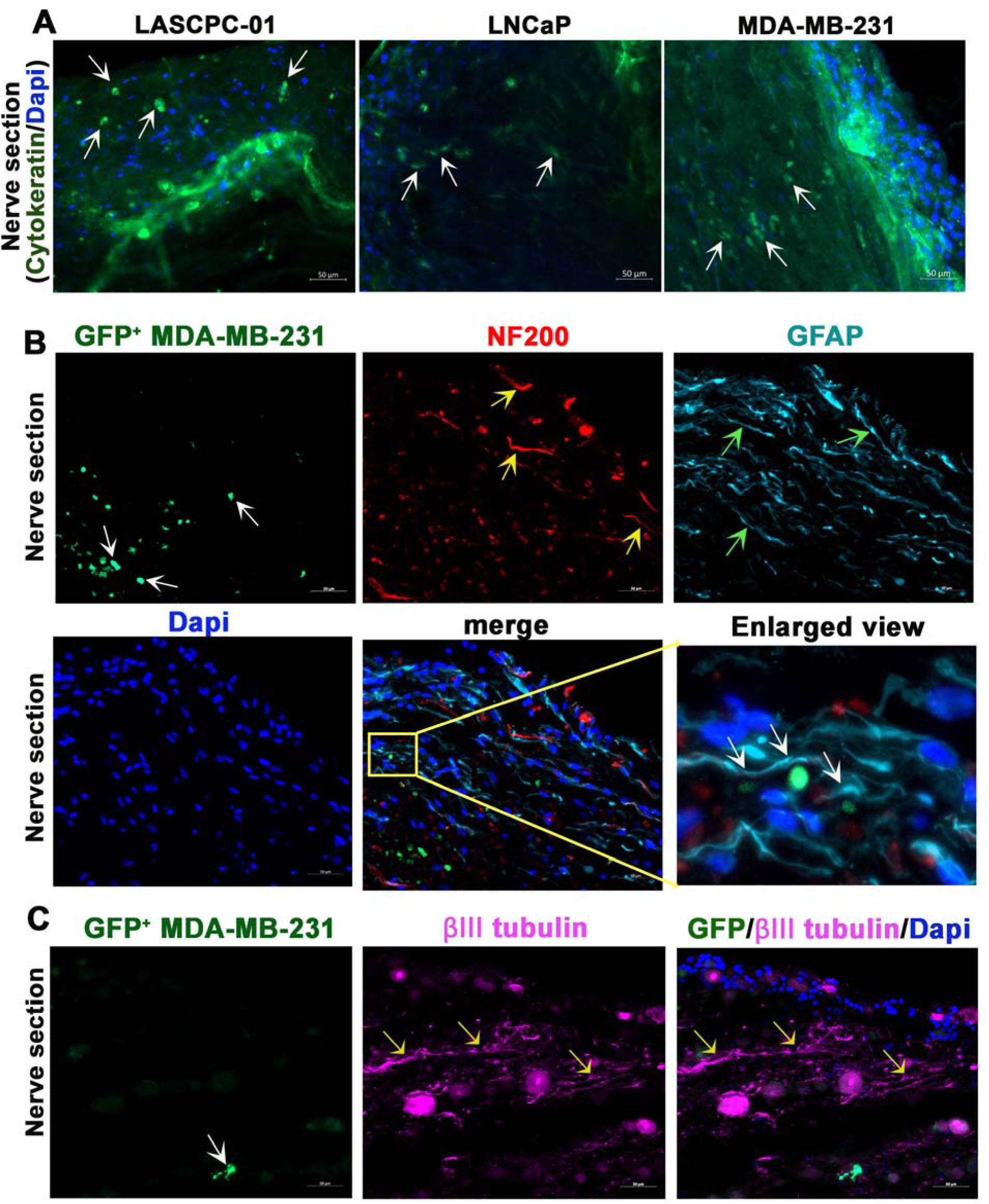
Cancer cells invade nerves in the *in vitro* PNI model: **(A)** Immunostaining of sections of DRG-nerve preparations isolated from the *in vitro* model (2-week time point) show the presence of cytokeratin positive cancer cells (yellow arrows) (scale bar, 50 µm). **(B; two rows)** Sections of DRG-nerve preparations isolated from the *in vitro* model (2-week time point) show the presence of GFP^+^ cancer cells (white arrows). Immunostaining of the sections show the presence of intact strands of NF200 positive axons (yellow arrows) and SCs (green arrows) (scale bar, 50 µm). Enlarged view of the merged image shows areas (white arrows) where cancer cells interact with SCs within the nerve. **(C)** Sections of DRG-nerve preparations isolated from the *in vitro* model (2-week time point) show the presence of GFP^+^ cancer cells (white arrows) and intact strands of βIII tubulin positive axons (yellow arrows) (scale bar, 50 µm).

In the DRGs, we found that PNI cells were preferentially distributed at the rim of neurons (**Figure 3A**). Satellite glial cells (SGCs), which are SC-like glial cells, naturally occupy the rim of injured neurons characterising neuron injury response. We found such rimming of neurons by SGCs in the *in vitro* PNI model indicating that most neurons in this model experience an injury response (**Figure 3B**). This injury response may be acquired during the tissue harvesting from experimental animals. Interestingly, the PNI cells did not overlap with SGCs’ locations in the DRGs, rather they preferentially located at neuronal rims free of SGCs (**Figure 3B**). Thus, the neurons interacting with PNI cells, instead of SGCs, may not experience an injury response. Alternatively, the interaction with cancer cells might blunt the neuron injury response. Regardless, the neuronal rimming and physical proximity of PNI cells to neurons suggest that neuron-derived factors may support the survival and integrity of PNI cells within the DRG milieu.

**Figure 3:**
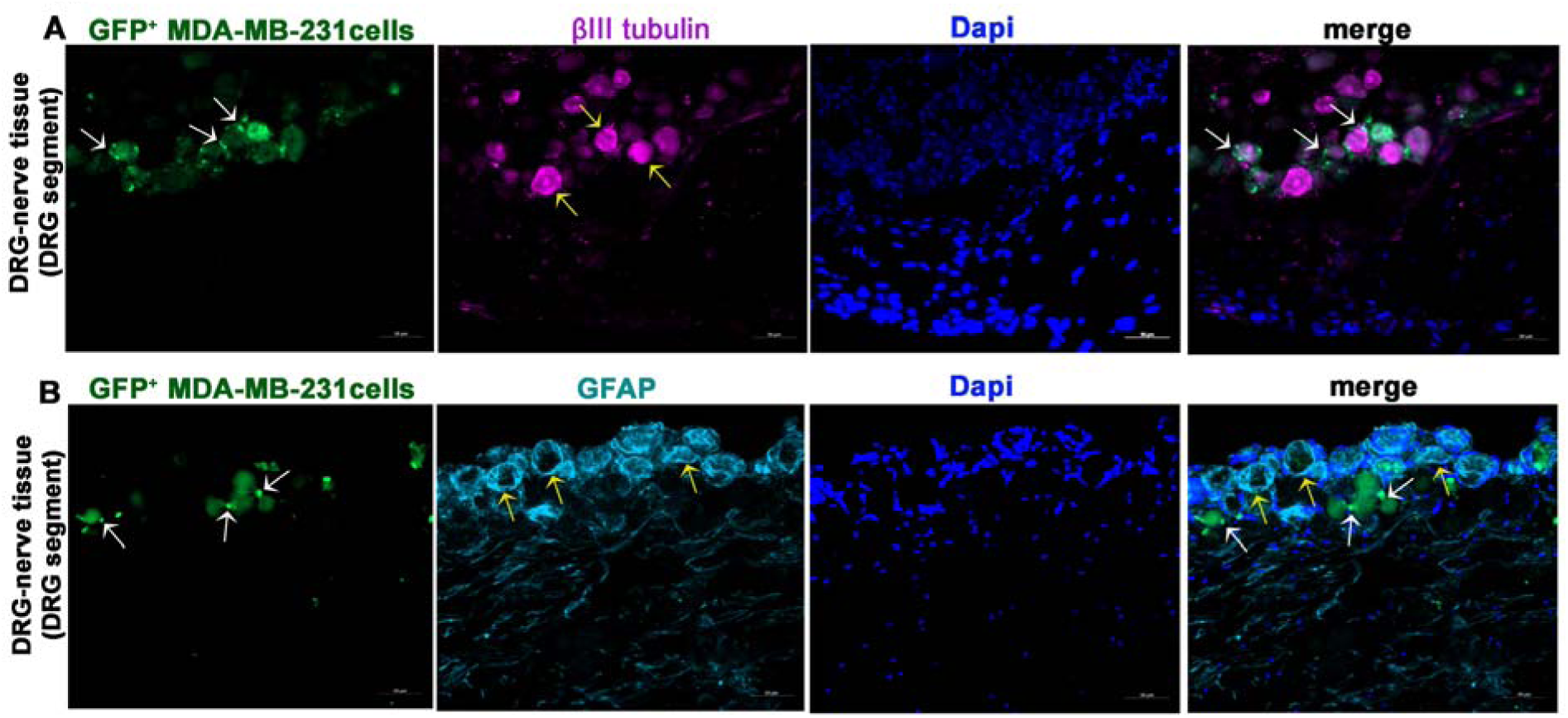
Cancer cells interact with DRG sensory neurons in the *in vitro* PNI model: **(A)** A section of DRG-nerve preparation from 2-week-old *in vitro* PNI model shows the presence of GFP^+^ cancer cells (white arrows) at the rim of neurons. The neurons are immunostained for βIII tubulin (yellow arrows) (scale bar, 50 µm). **(B)** A section of DRG-nerve preparation from 2-week-old *in vitro* PNI model shows no physical overlap of cancer cells (white arrows) with SGCs (yellow arrows) in the DRG (scale bar, 50 µm).

### Basic and therapeutic studies related to PNI using the *in vitro* PNI model

Previous research showed that PNI-competent prostate cancer cells have a lower apoptotic index, and thus, possess survival advantages^13^. Similarly, autonomic neurosignaling, especially signaling driven by the neurotransmitters acetylcholine (ACh) and norepinephrine (NE), has been shown to promote prostate cancer progression^14–16^. Parasympathetic (cholinergic) neurosignaling was also shown to promote PNI ^17, 18^. Therefore, we examined the influence of ACh and NE on PNI incidence using the *in vitro* model. We also examined the influence of ACh and NE on cisplatin-induced toxicity of PNI cells. For this, *in vitro* co-cultures of DRG-nerve and GFP^+^ LASCPC-01 cells were supplemented with ACh (10 µM) or NE (10 µM) every three days for two weeks. Then after, the cultures were treated with 50 µM cisplatin for 48 h. The PNI incidence was quantified by manual counting of GFP^+^ cancer cells in the DRG-nerve sections. We found that both NE and ACh promoted PNI incidence. Although cisplatin treatment reduced the number of PNI cells compared to control (saline) treatment, the effect was not significant, indicating that PNI cells are at least partially protected from cytotoxic treatments (**Figure 4A, B**).

**Figure 4:**
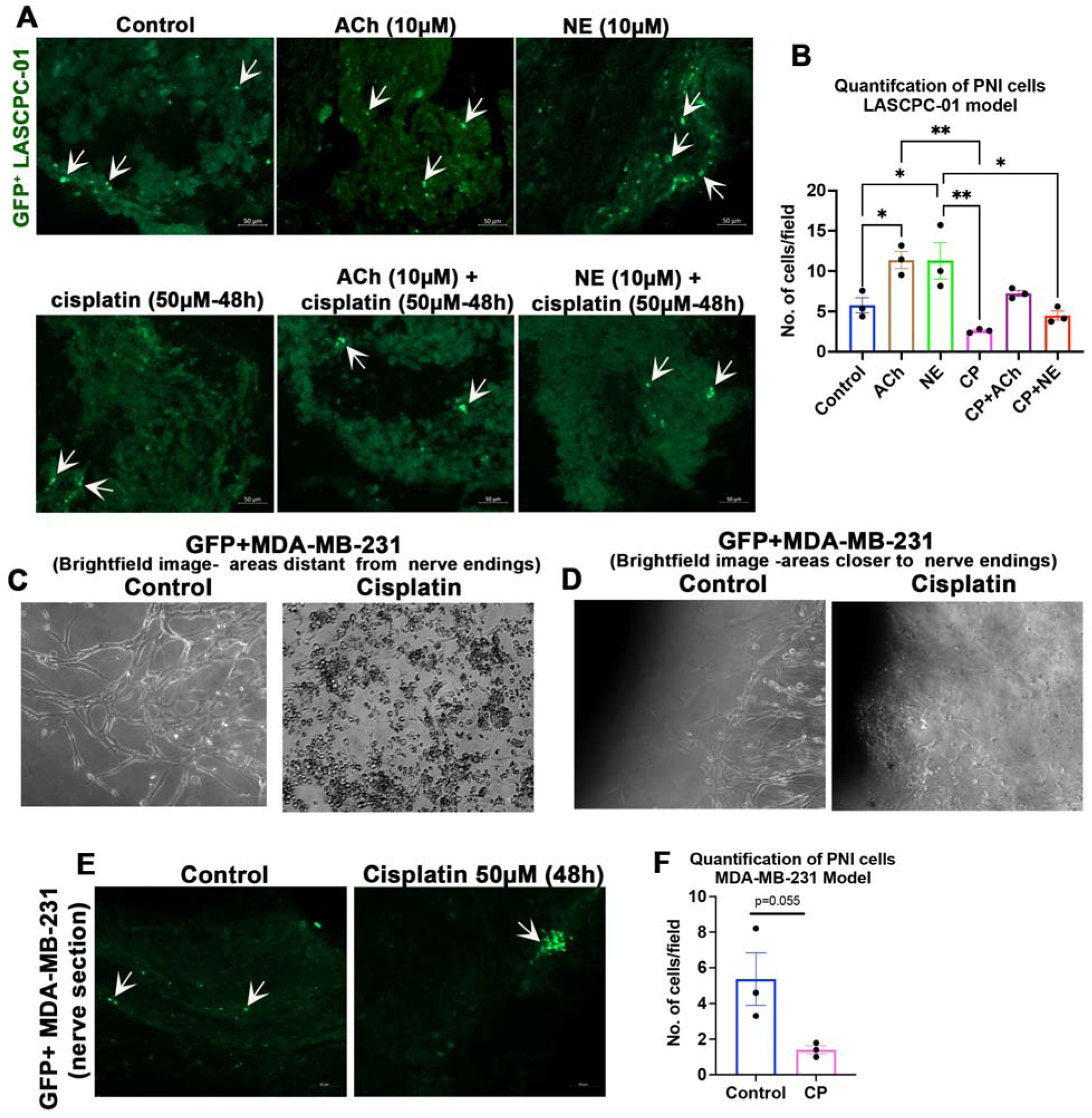
Autonomic signaling promotes PNI and PNI cells have survival advantages against chemotherapy treatment: **(A)** Sections of 16-day-old DRG-nerve preparations subjected to the indicated treatments in the *in vitro* model show healthy PNI-competent prostate cancer cells (white arrows) (scale bar, 50 µm). **(B)** Quantification of ‘A’ (n=3, One-Way ANOVA with Tukey’s multiple comparisons test, *p<0.05; **p<0.01). **(C)** Bright field images show remarkable cell death of MDA-MB-231 cells in areas distant from the nerve endings in the *in vitro* PNI model after cisplatin (50 µM) treatment for 48 h (scale bar, 100 µm). (D) Bright field images show remarkable cell death of MDA-MB-231 cells in areas closer to the nerve endings in the *in vitro* PNI model after cisplatin (50µM) treatment for 48 h (scale bar, 100 µm). **(E)** Sections of 16-day-old DRG-nerve preparations subjected to control (saline) and cisplatin treatment in the *in vitro* model show healthy PNI-competent breast cancer cells (white arrows) (scale bar, 50 µm). (F) Quantification of PNI cells in sections of DRG-nerve preparations following cisplatin (50 µM) treatment every three days for 14 days (n=3, Student’s t-test).

We next performed similar survival experiments using GFP^+^ MDA-MB-231 cells. Brightfield microscopy revealed remarkable cell death exterior to nerves after cisplatin treatment indicating the vulnerability of cells exterior to the nerves to cisplatin treatment (**Figure 4C, D**). In contrast, sections of nerves showed intact GFP^+^ MDA-MB-231 cells in both control and cisplatin groups indicating protection of PNI cells from cisplatin (**Figure 4E**). Since cancer patients receive multiple doses of chemotherapy as part of their treatment regimen, we next evaluated the effect of repeated treatments of cisplatin on PNI incidence. For this, the DRG-nerve-cancer cell co-cultures were treated with 50 µM cisplatin every three days for two weeks. This continuous treatment paradigm too, although eliminated PNI cells, did not elicit a significant effect (**Figure 4F**). Overall, our results indicate that PNI cells have survival advantages, and most importantly, the experiments described above demonstrate the utility of the newly characterized PNI model in studying the fundamentals of PNI and screening potential anti-PNI therapies.

### Molecular characterization of PNI-competent breast cancer cells using proteomics studies

Molecular characterization of PNI cells should facilitate the development of theranostics approaches, enabling accurate diagnosis and treatment of PNI, to eventually reduce PNI-dependent recurrence and metastasis. Thus, we characterized the molecular signature of PNI-competent breast cancer cells using the newly established PNI model. For this, we sorted PNI-competent GFP^+^ MDA-MB-231 cells from two-week old PNI model using a flow cytometer. PNI cells from about 50 experimental preparations were pooled to generate a single replica. The sorted PNI cells were then subjected to whole cell proteomics analysis. The PNI-incompetent cells harvested from areas in the Cultrex matrix physically away from the DRG-nerve preparations were used as the control. Two replicas, each for the test (purified PNI cells) and control (PNI-incompetent cells), were used for the proteomics analysis.

The differentially expressed proteins (≥ 2-fold; p<0.05) identified in PNI-competent cells from the proteomics analysis are listed in **Table S1**. Of the 382 differentially expressed proteins (after eliminating keratin, type II cytoskeletal 2 epidermal and keratin, type I cytoskeletal 9, which are common contaminants as described in the common repository of adventitious proteins (cRAP)), 377 proteins were significantly downregulated in PNI cells compared to the control. Panther Gene Ontology (GO) analysis showed that molecules associated with binding (integrin, cadherin, Rho GTPase), catalytic activity (molecules associated with glucose metabolism), and structural molecular activity (integrin, cadherin, Rho GTPase) were predominantly downregulated in PNI cells. Most importantly, the downregulation of catalytic proteins related to glucose metabolism suggests a low metabolic profile for PNI cells, indicating that PNI cells may enter an inactive state (**Figure 5B & Figure S2**). Five proteins, including the cilia and planar cell polarity associated protein 1 (CPLANE1), hemopexin, SANT and BTB Domain Regulator of Class Switch Recombination (SANBR)^19^, and ATM, the DNA repair and cell cycle check-point kinase^20, 21^, were significantly upregulated in PNI cells. The functions of CPLANE1 are primarily associated with cilia biogenesis, actin organization, and cell polarity and these functions can potentially promote PNI^22^. STRING analysis predicted interactions of hemopexin with potential PNI modulators, including matrix metalloproteinases (MMPs), suggesting a prominent role for hemopexin in modulating PNI **(Figure 5C).** The upregulation of SANBR and ATM suggests potential DNA rearrangements in PNI cells. Overall, the global downregulation of proteins, including the ones associated with metabolic activities, coupled with the induction of proteins associated with DNA modifications, are suggestive of a dormant status of PNI cells within the nerve milieu. Additionally, we found no co-localization of PNI cells and the cell proliferation marker Ki67 within nerves, further supporting the argument that PNI cells may enter dormancy (**Figure S3**). However, additional systematic studies are warranted to confirm the specific phenotype acquired by the PNI cells within the nerve milieu.

**Figure 5:**
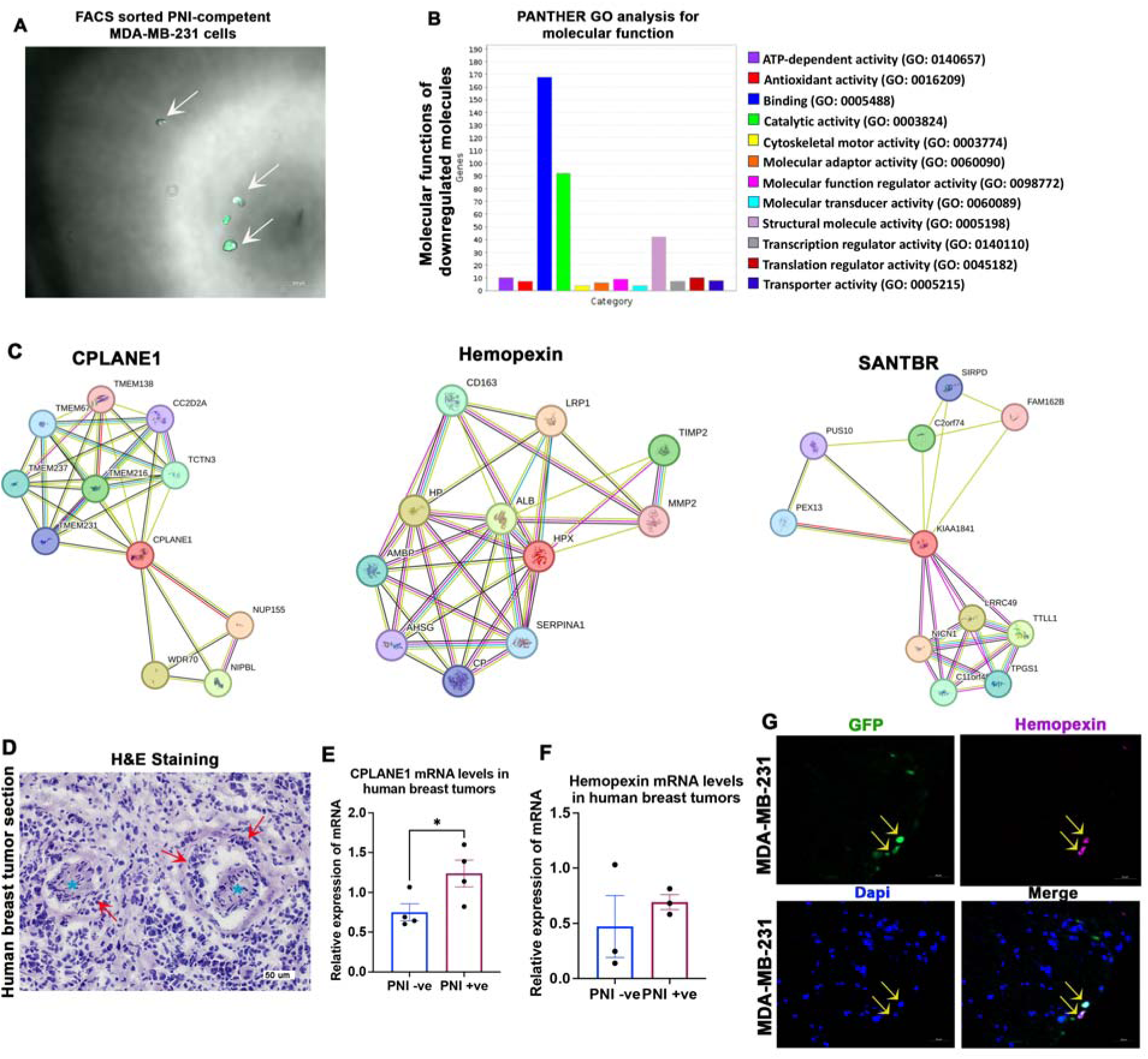
Proteomics characterization of PNI cells. **(A)** Combined brightfield and fluorescence image of a droplet of FACS-sorted PNI cells shows the purity of the isolated cells (white arrows). **(B)** PANTHER GO analysis shows the enriched molecular functions of molecules downregulated in PNI cells. **(C)** STRING analysis shows protein interaction network of CPLANE1, hemopexin and SANBR. **(D)** A representative image of a hematoxylin and eosin (H&E) stained human breast tumor sample showing PNI (red arrows; the cyan asterisks indicate nerves) (scale bar, 50 µm). **(E)** CPLANE mRNA levels in PNI^+^ and PNI^−^ human breast tumor samples (n=4; Student’s ‘t’ test, *p<0.05). **(F)** Hemopexin mRNA levels in PNI^+^ and PNI^−^ human breast tumor samples (n=3; Student’s ‘t’ test). **(G)** Representative image of a section of DRG-nerve preparation from 2-week-old *in vitro* PNI model shows the expression of hemopexin in PNI cells (scale bar, 50 µm).

### CPLANE1 and hemopexin may serve as molecular markers of PNI cells

Given the known role of CPLANE1 in cell polarity and hemopexin’s potential connection with MMPs, we postulated that they contribute to PNI. To verify this, we validated their expression in human PNI cells. First, we examined PNI occurrence in a total of 98 H&E-stained human breast tumor samples and found PNI in six samples, accounting for an overall PNI incidence of ∼ 6% (**Figure 5D)**. Among the six PNI positive (PNI^+^) cases, four patients developed metastasis either in the brain or bone **(Table S2),** accounting for more than 60% of PNI^+^ cases resulting in metastasis. We then examined the mRNA levels of CPLANE1 and hemopexin in a small cohort of PNI^+^ and PNI^−^ tumor samples and found significant upregulation of CPLANE1 in PNI^+^ cases validating the proteomics results (**Figure 5E)**. Although not significant, we also found a trend in the upregulation of hemopexin mRNA in PNI^+^ samples (**Figure 5F).** Next, we examined the expression of hemopexin in our *in vitro* model using immunostaining and found that it is specifically expressed in PNI cells within the nerve with no expression observed in other cell types, including axons and SCs. This finding indicate that hemopexin may serve as a unique marker for PNI cells, enabling the development of targeted theranostics approaches **(Figure 5F).**

## Discussion

Numerous studies have reported that PNI is a risk factor for cancer metastasis and recurrence, linking PNI incidence with poor prognosis of cancers^1–4^. PNI involves the intricate interaction between nerves and cancer cells enabling cancer cells’ invasion into internal layers of peripheral nerves. However, to our knowledge, no *in vitro* PNI model demonstrate the interaction of cancer cells with axons, neurons, and SCs within nerves, and thus, fail to recapitulate the complex nerve-cancer cell interaction. This was indeed a limitation in advancing the knowledge about PNI biology and screening potential therapies. Here, we characterized an *in vitro* model for PNI, recapitulating the complex nerve-cancer cell interaction by employing intact DRG-nerve preparations. Using this model, we demonstrated the induction of natural PNI *in vitro*, revealed the distribution of PNI cells within the nerve milieu, and identified their interactions with axons, SCs and neurons. To our knowledge, this is the first fully characterized *in vitro* model that closely mimics natural PNI.

*In vitro* preparations of intact nerves are challenging to develop due to the immediate degeneration of axons upon losing contact with neurons^23^. Axons, the conducting fibres of nerves, secrete growth factors to support their companion SCs^23^. Conversely, the SCs in contact with axons also secrete neurotrophic factors to support the homeostasis of axons^23^. It is thus critical to preserve this reciprocal growth exchange environment in *in vitro* nerve preparations when modeling natural PNI *in vitro* as these growth factors may also contribute to PNI. The *in vitro* PNI model characterized in this work involves a nerve with its axons intactly connected to sensory neurons housed in the DRGs. As such, this model preserves axon integrity by maintaining their intact connection with neuron cell bodies. The integrity of axons in this model was further confirmed by visualizing healthy axons in the tissue preparation after staining them with the pan-neuronal marker βIII tubulin. However, we found partial loss of NF200^+^ axons in this model. NF200 is a cytoskeletal protein expressed preferentially in large diameter axons. NF200, especially dephosphorylated NF200, loses its expression after traumatic nerve injuries^24^. We believe that the loss of NF200 in the *in vitro* PNI model may reflect an injury response following the harvesting of DRG-nerves from animals. This nerve injury response in the *in vitro* model can be acceptable as tumor infiltrated nerves are often compressed or damaged by growing tumors, potentially initiating a similar injury response.

The *in vitro* model presented here showed that SCs emerging from nerve endings provide tropism signals to cancer cells for PNI. This model also showed that PNI cells interact with SCs within nerves. SCs have previously been shown to promote PNI^25^, and hence, the cancer cell-SCs crosstalk observed in this model substantiates the occurrence of natural PNI. At the same time, molecular cues emanating from axons may also contribute to PNI. We believe that this model provides a valuable platform for exploring these molecular players, with potential application in future theranostics development.

We used adenoviral particles to generate GFP^+^ cancer cells for tracking PNI cells within nerves. As this system does not generate stable cell clones, our experiments might have not captured all PNI cells during imaging. Despite this limitation, our experiments successfully identified PNI incidence, characterized PNI cells’ distribution, and revealed their crosstalk with axons and SCs within the nerve milieu. Additionally, the *in vitro* model established here enabled us to characterize the proteomics profile of PNI cells. As such, we do not believe that the use of an adenoviral system resulted in the loss of any key information. However, future studies employing cancer cells with stable markers will provide additional clarity.

Although we used breast and prostate cancer cells to characterize the model, the proteomics characterization was done using PNI-competent breast cancer cells. Therefore, the molecular signature of PNI cells identified in this study may not be directly applicable to PNI-competent prostate cancer cells. While our *in vitro* model demonstrated that breast and prostate cancer cells both follow a similar pattern of distribution and engagement within nerves, the PNI-related molecular players intrinsic to these cells may be different. We found that CPLANE1 is upregulated in PNI-competent breast cancer cells. CPLANE1 promotes ciliogenesis and cell polarity, both of which are favorable for PNI^22^. Ciliogenesis is also linked with quiescence state, suggesting that PNI cells may enter dormancy within the nerve milieu^26^. Additionally, the global downregulation of proteins in PNI-competent breast cancer cells also suggests that PNI cells may enter dormancy within the nerve environment. The upregulation of SANBR-1 and ATM in PNI cells may indicate potential DNA rearrangements, favoring remarkable molecular reprogramming and related phonotypic reprogramming directed towards dormancy. Interestingly, we observed induction of hemopexin in PNI-competent breast cancer cells. Hemopexin has recently gained attention for its role in promoting cancer progression and metastasis^27^. However, a recent study also showed that hemopexin suppresses cancer cell dissemination by sequestering heme in the TME^28^. The unique presence of hemopexin in PNI cells is thus interesting and suggests that hemopexin may facilitate the retention of dormant PNI cells within the nerve milieu. However, further studies are needed to draw definite conclusions. Overall, based on our observations, we speculate that PNI cells may enter dormancy within the nerve milieu. In clinical recurrence, the dormant PNI cells may undergo reactivation when signals favorable for their revival build up within nerves. However, more systematic studies are required to support this argument.

Like all *in vitro* experimental models, this model is also not devoid of limitations. The model requires careful isolation of DRG-nerves from adult rats. Overstretching of nerves during isolation may damage axons entirely and affect the reproducibility of the model. The method described in this manuscript used adenoviral system to generate GFP^+^ cancer cells (transient expression of GFP), with a limitation of missing the capturing of GFP-lost PNI cells during imaging. The future use of cancer cells stably expressing GFP will ensure the capturing of all PNI cells. Another limitation is the poor yield of PNI cells in this natural invasion model for molecular biology studies. Pooling PNI cells from several replicas, as done in this work, will resolve the issue.

The distinguishing features of the currently available *in vitro* models of PNI and the explant model characterized in this work are summarized in **Figure 6**.

**Figure 6:**
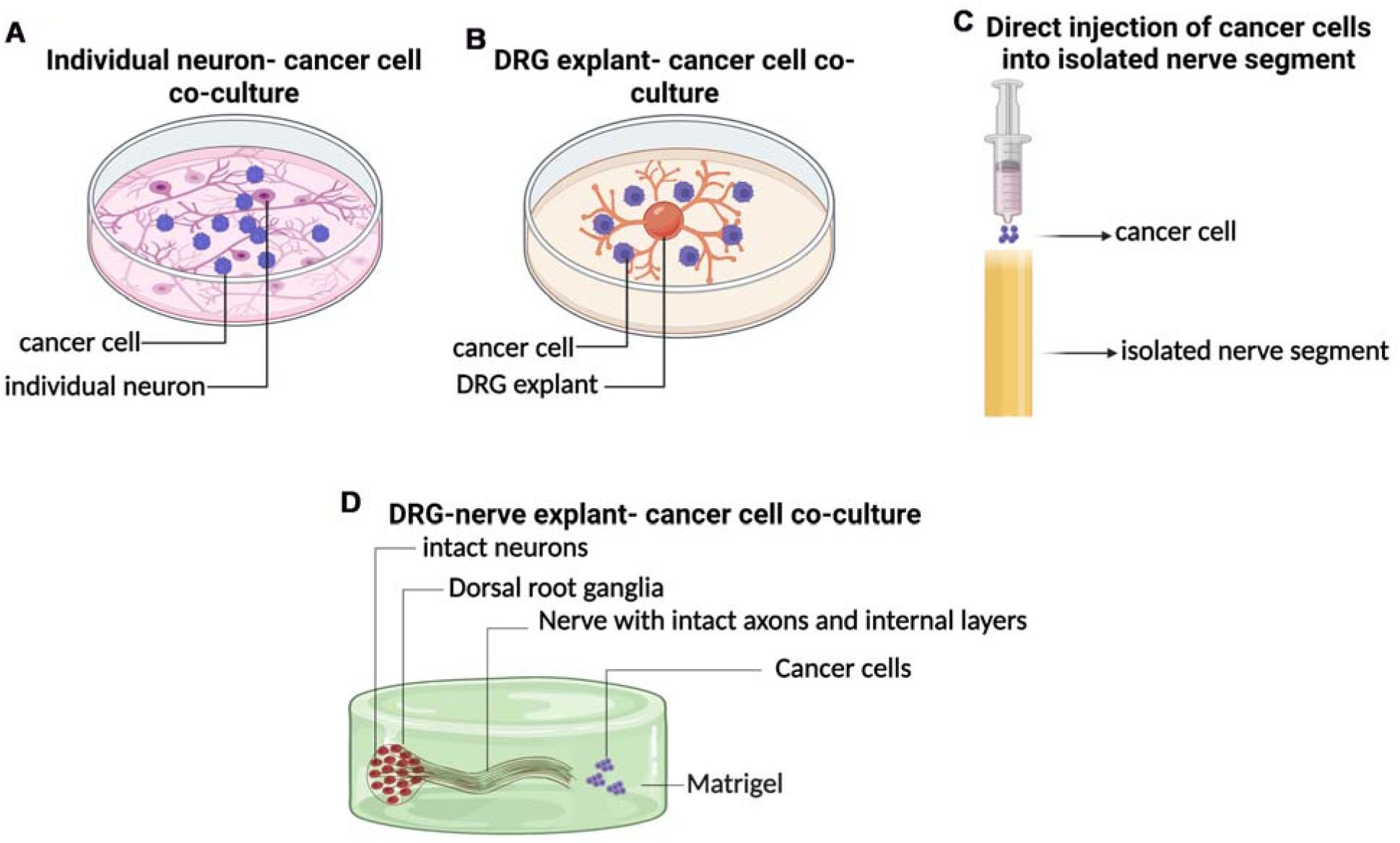
Distinguishing features of the available and newly established *in vitro* PNI models: **(A**) *Co-culture of individual neurons and cancer cells:* This preparation has no nerve scaffold, and is thus devoid of the internal layers of the nerve, which are critical for PNI to occur. **(B)** *Co-culture of DRG explant and cancer cells:* This preparation has no nerve scaffold, and is thus devoid of the internal layers of the nerve, which are critical for PNI to occur. **(C)** *Forced injection of cancer cells into isolated nerve scaffold:* This preparation although retains a nerve scaffold, will not have intact axons (the fundamental units of nerve). Note that axons degenerate immediately after detaching from neurons (cell bodies), and hence, a cut nerve without neuron cell bodies will not represent a physiological model as there will be only degenerated axons within the nerve scaffold. **(D)** *Co-culture of DRG-nerve explant and cancer cells (the in vitro model established in this work):* This model has a nerve segment intactly connected to neuron cell bodies. Thus, this preparation maintains intact axons, allowing natural interaction of the invaded cancer cells with axons, neurons and SCs. This model also maintains intact nerve layers for natural PNI to occur.

## Conclusion

In conclusion, we characterized and established an *in vitro* PNI model for studying the fundamentals of PNI and screening potential anti-PNI therapies. We have also characterized the molecular profile of PNI-competent breast cancer cells and showed that CPLANE1 and hemopexin are potential markers for PNI-competent breast cancer cells. Additional studies are warranted to understand the functional roles of CPLANE-1 and hemopexin in PNI and to explore the roles of other players, including SANBR-1.

## Author contributions

NJ performed experiments, analyzed data, and contributed to preparing the initial and revised versions of the manuscript. SS, IS, SM, PC, BS, and VB contributed to data generation and analysis. SA and HR contributed to data analysis and edited the manuscript. GSK provided the mass spectrometry instrumentation, analyzed data, and contributed to preparing the initial and revised versions of the manuscript. AK conceived the concept, analyzed the data, prepared the initial and revised versions of the manuscript, and finalized the manuscript.

## Ethics statement

The purchase of human breast tumor samples and the use of associated de-identified clinical data have been approved by the Biomedical Research Ethics Board of the University of Saskatchewan. Adult rats used for harvesting the DRG-nerve preparations have been approved by the Animal Research Ethics Board at the University of Saskatchewan.

## Acknowledgements

This work was supported by the Establishment Grant from the Saskatchewan Health Research Foundation (SHRF) to AK. This work was also supported by the Operating Grants from the Breast Cancer Society of Canada and the Prostate Cancer Fight Foundation and Ride for Dad to AK. We thank the Alberta Cancer Research Biobank (ACRB) for providing us with the human cancer samples.

## Data availability

Data will be made available upon reasonable request.

