## Supplementary figures for "Defining the cellular and molecular features of nerve-invaded cancer cells using a newly characterized experimental model"

**Supplemental information**

**Figure S1:** (**A**) Immunostaining shows the distribution of NF200^+^ and βIII tubulin^+^ axons in a normal sciatic nerve (scale bar, 100µm). (**B**) A section of DRG-nerve preparation isolated from the *in vitro* model (2-week time point) shows the presence of GFP^+^ prostate cancer cells (white arrows). The white arrows in the merged image shows areas where cancer cells interact with SCs within the nerve (scale bar, 50µm).

**Figure S2:** Pie charts show that molecules associated with integrin, cadherin, and cytoskeletal signaling and pyruvate and glucose metabolism are downregulated in PNI cells.

**Figure S3:** Immunostaining of 14-day PNI model shows no co-localization of Ki67 (cell proliferation marker; white arrows) and PNI cells (green; yellow arrows).

**Table S1:** Excel file containing differentially regulated list of proteins from the proteomics data analysis.

**Table S2:** Excel file containing the clinical parameters associated with the human cancer samples used for the study.

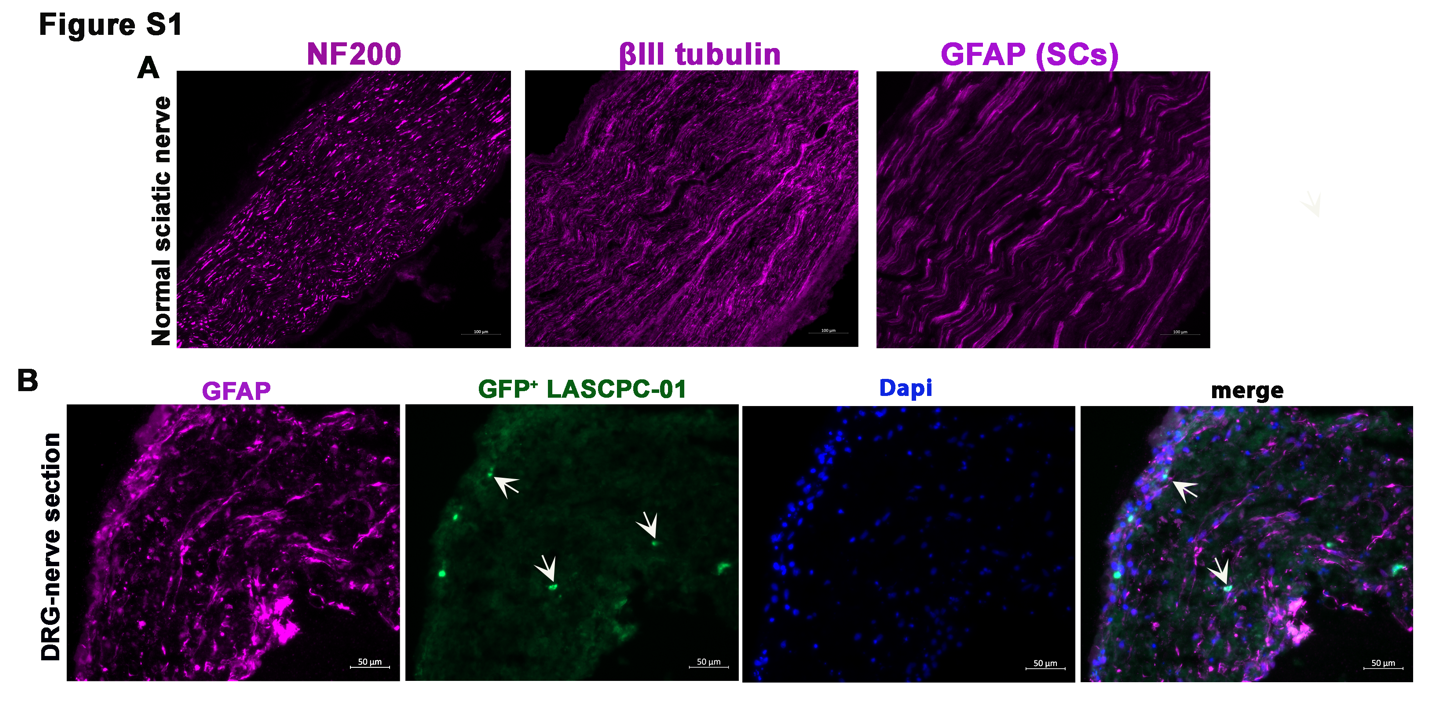

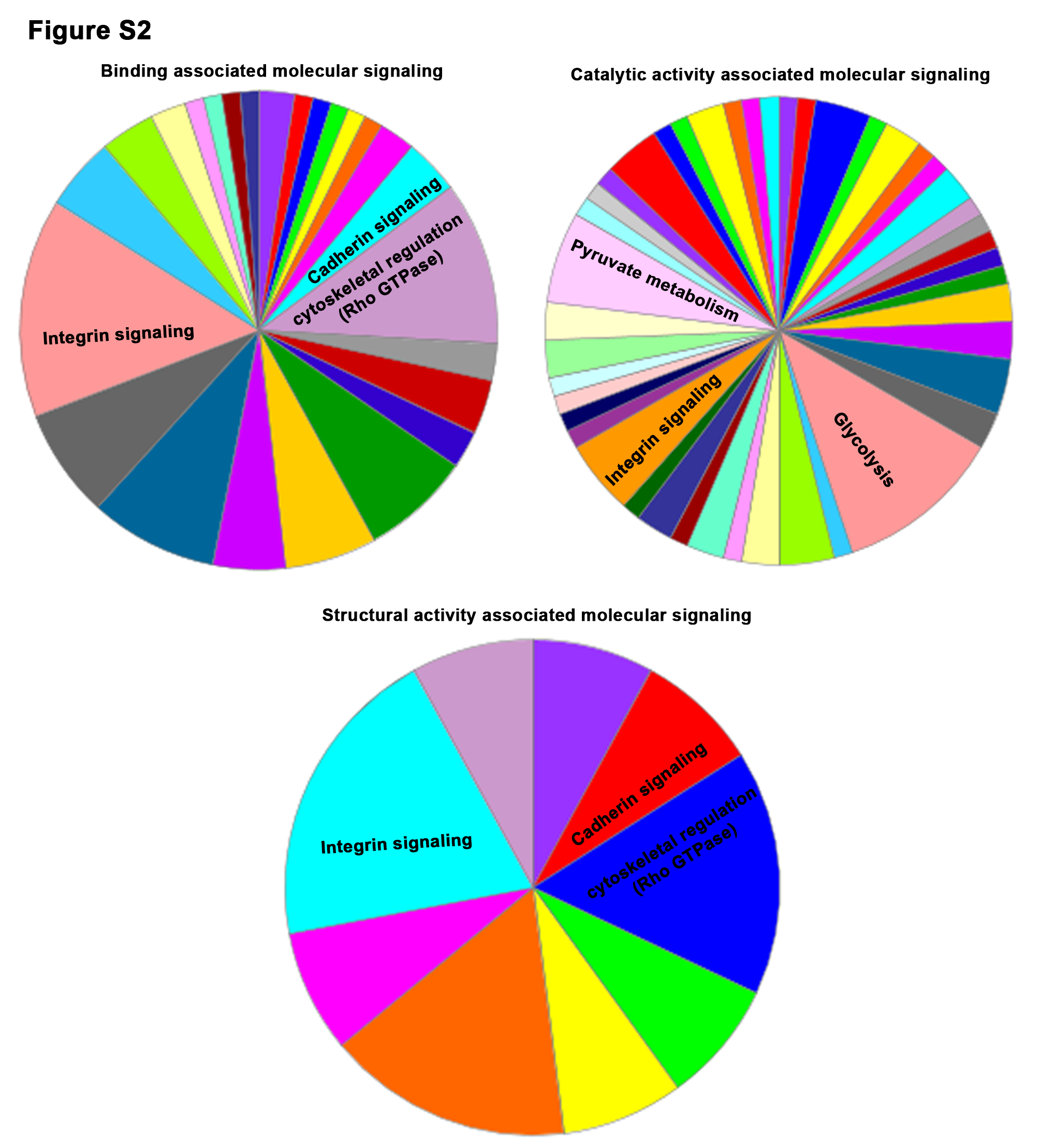

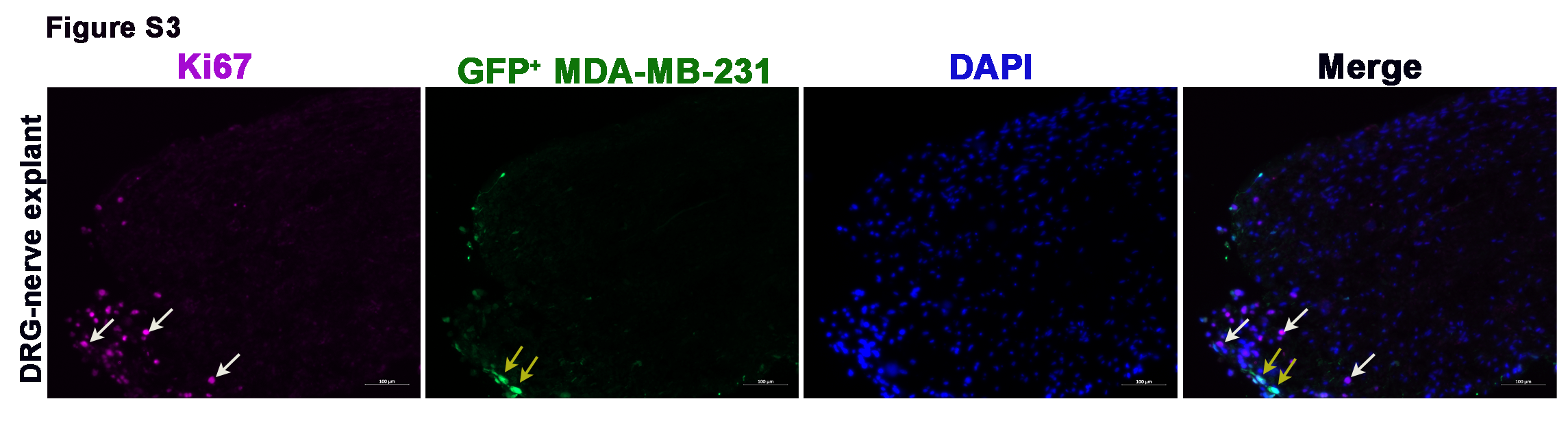
